# Predicting microbial relative growth in a mixed culture from growth curve data

**DOI:** 10.1101/022640

**Authors:** Yoav Ram, Eynat Dellus-Gur, Maayan Bibi, Uri Obolski, Judith Berman, Lilach Hadany

**Affiliations:** Dept. Molecular Biology and Ecology of Plants, Tel Aviv University, Tel Aviv 69978, Israel.; Dept. of Molecular Microbiology and Biotechnology, Tel Aviv University, Tel Aviv 69978, Israel.

**Author notes:** Current address: Dept. of Biology, Stanford University, Stanford, CA 94305. Current address: Dept. of Zoology, University of Oxford, Oxford, UK.

**Keywords:** competition models, growth models, fitness, experimental evolution, microbial evolution, microbial ecology

## Abstract

Estimates of microbial fitness from growth curves are inaccurate. Rather, competition experiments are necessary for accurate estimation. But competition experiments require unique markers and are difficult to perform with isolates derived from a common ancestor or non-model organisms. Here we describe a new approach for predicting relative growth of microbes in a mixed culture utilizing mono- and mixed culture growth curve data. We validated this approach using growth curve and competition experiments with *E. coli*. Our approach provides an effective way to predict growth in a mixed culture and infer relative fitness. Furthermore, by integrating several growth phases, it provides an ecological interpretation for microbial fitness.

## Introduction

Microbial fitness is usually defined as the relative growth of different microbial strains or species in a mixed culture^1^. Pairwise competition experiments can provide accurate estimates of relative growth and fitness^2^, but they are laborious and expansive, especially in non-model organisms. Instead, growth curves are commonly used to estimate fitness of individual microbial isolates, despite clear evidence that they provide an insufficient alternative^3,4^.

Growth curves describe the density of cell populations in liquid culture over time and are usually acquired by measuring the optical density (OD) of one or more cell populations. The simplest way to infer fitness from growth curves is to estimate the growth rate during the exponential growth phase by inferring the slope of the log of the growth curve^5^ (see example in Figure 1). Indeed, the growth rate is often used as a proxy of the selection coefficient, *s*, which is the standard measure of relative fitness in population genetics^1,6^. However, exponential growth rates do not capture the dynamics of other phases of a typical growth curve, such as the length of lag phase and the cell density at stationary phase^7^ (Figure 1A). Moreover, the maximal specific growth rate is not typical for the entire growth curve (Figure 1B). Thus, it is not surprising that growth rates are often poor estimators of relative fitness^3,4^.

**Figure 1.**
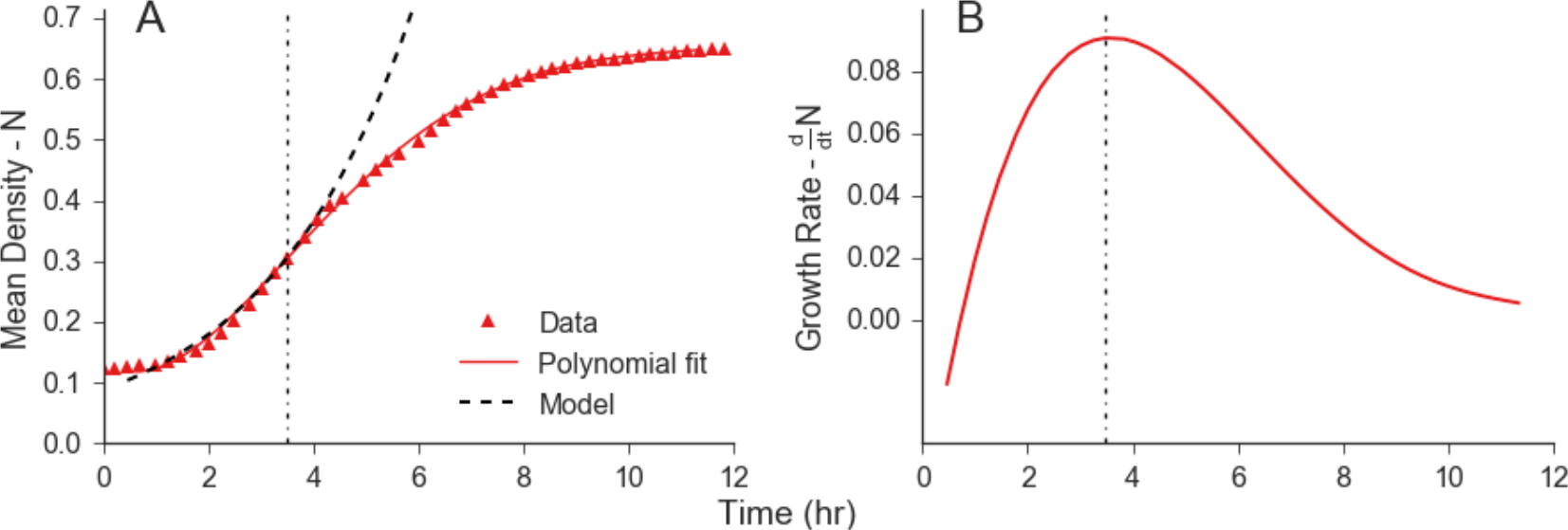
Common approach for analyzing growth curve data using an exponential model. Growth rates are commonly estimated from growth curves data by taking the log of the growth curve and performing linear regression around the time of maximum growth (see Materials and Methods for specific details). Implicitly, this is equivalent to fitting an exponential growth model *N(t)=N_0_e^rt^* to the growth curve. **(A)** The red markers represent *N(t)* the mean cell density in 22 growth curves. The solid red line represents a smooth line through the points (e.g. by fitting a polynomial). The dashed black line represents the exponential model *N_0_e^rt^* fitted to the data, with *r=0.27* and *N_0_=0.058*. The dotted vertical line denotes *t_max_*. **(B)** The red solid curve shows *dN/dt,* the derivative of the mean density (calculated as the derivative of the red line in A). The dotted vertical line denotes *t_max_*. Data in this figure corresponds to the growth of the red strain in experiment A (red markers in Figure 2A).

In contrast, competition experiments can infer relative fitness in a manner that accounts for all growth phases^8^. In pairwise competition experiments, two strains are grown together in a mixed culture: a reference strain and a strain of interest. The frequency of each strain in the mixed culture is measured during the course of the experiment using specific markers, for example, by counting colonies formed by drug resistant or auxotrophic strains^8^, by monitoring fluorescent markers with flow cytometry^2^, or counting DNA barcodes reads using deep sequencing^9,10^. The selection coefficient of the strains of interest can then be estimated from changes in their frequencies during the competition experiments. These methods can infer relative fitness with high precision^2^, as they directly estimate fitness from changes in strain frequencies over time.

However, competition experiments are more laborious and expensive than growth curve experiments, requiring the development of genetic or phenotypic assays (see Concepción-Acevedo et al.^3^ and references therein). Moreover, competition experiments are often impractical in non-model organisms. Therefore, many investigators prefer to use proxies of fitness such as growth rates. Even when competition experiments are a plausible approach (for example, in microbial lineages with established markers^8^), methods for interpreting and understanding how differences in growth contribute to differences in fitness are lacking. Such differences have a crucial impact on our understanding of microbial fitness and the composition of microbial populations and communities.

Here we present a new computational approach which provides a predictive and descriptive framework for estimating growth parameters from growth dynamics and predicting relative growth in mixed cultures.

## Results

Our approach consists of three stages: (a) fitting growth models to monoculture growth curve data, (b) fitting competition models to mixed culture growth curve data, and (c) predicting relative growth in a mixed culture using the estimated growth and competition parameters. To test our approach, we measured growth of two *E. coli* strains in mono- and mixed culture over time and used our new approach to predict the relative frequencies of both strains in the mixed culture. We then compared these predictions to empirical measurements of strain frequencies.

## Experimental design

Before proceeding to describe our new approach in detail, we briefly describe the experiments we used to test it, as results of these experiments will be presented in the following sections. We performed growth curve and competition experiments with two different sets of *E. coli* strains marked with green and red fluorescent proteins (GFP and RFP, respectively). In each experiment, 32 replicate monocultures of the GFP strain, 30 replicate monocultures of the RFP strain alone, and 32 replicate mixed cultures containing the GFP and RFP strains together, were grown in a 96-well plate, under the same experimental conditions. In experiments A and B we used *E. coli* strains DH5α-GFP and TG1-RFP; in experiment C we used *E. coli* strains JM109-GFP and MG1655-Δfnr-RFP. The optical density of each culture was measured every 15 minutes using an automatic plate reader for at least 7 hours (Figure 2A-C). Samples were collected from the mixed culture every hour for the first 7-8 hours, and the relative frequencies of the two strains were measured by flow cytometry. See

Materials and Methods for additional details.

## a. Estimate growth parameters

### Growth model

The Baranyi-Roberts model^11^ can be used to model growth composed of several phases: lag phase, exponential phase, deceleration phase, and stationary phase^5^. The model implicitly assumes that growth accelerates as cells adjust to new growth conditions, then decelerates as resources become scarce, and finally halts when resources are depleted^12^. The model is described by the following ordinary differential equation (see eqs. 1c, 3a, and 5a in Baranyi and Roberts, 1994^11^; for a derivation of eq. 1 and further details, see Supporting Txt 1):

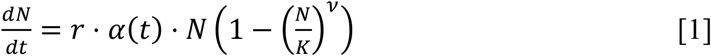

where *t* is time, *N* = *N*(*t*) is the cell density at time *t, r* is the specific growth rate in low density, *K* is the maximum cell density, *ν* is a deceleration parameter, and *α*(*t*) is the adjustment function, which describes the fraction of the population that has adjusted to the new growth conditions by time *t* (*α*(*t*) ≤ 1). In microbial experiments, an overnight liquid culture of microorganisms that has reached stationary phase is typically diluted into fresh media. Following dilution, cells enter lag phase until they adjust to the new growth conditions. We chose the specific adjustment function suggested by Baranyi and Roberts^11^, which is both computationally convenient and biologically interpretable: 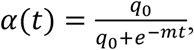 where *q*_0_ characterizes the physiological state of the initial population, and *m* is the rate at which the physiological state adjusts to the new growth conditions.

The Baranyi-Roberts differential equation (eq. 1) has a closed form solution:

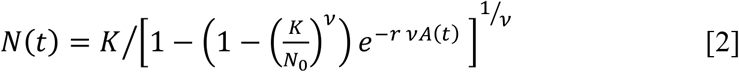

where *N*_0_ = *N*(0) is the initial population density. For a derivation of eq. 2 from eq. 1, see Supporting Text 1.

### Model fitting

We estimated the growth model parameters by fitting the model (eq. 2) to the monoculture growth curve data of each strain. The best-fit models (black lines) and experimental data (markers) are shown in Figure 2; see Table S1 for the estimated growth parameters. From these best-fit models we also estimated the maximum specific growth rate 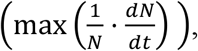 the minimal specific doubling time (minimal time required for cell density to double), and the lag duration; see Table 1. The strains differ in their growth parameters: for example, in experiment A (Figure 2A), the red strain grows 41% faster than the green strain, has 23% higher maximum density, and a 60% shorter lag phase.

**Figure 2.**
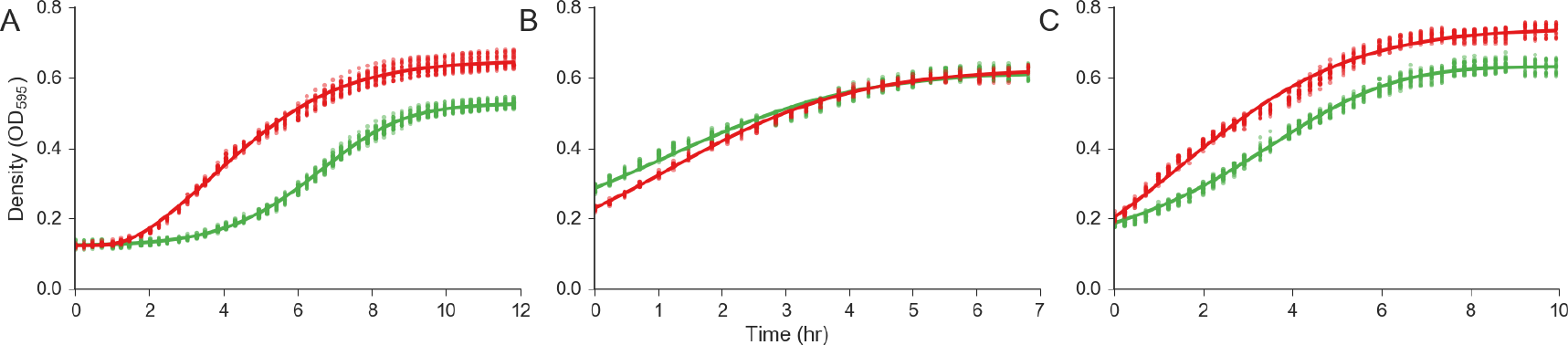
Fitting the growth model to data from three growth curve experiments with *E. coli*. Colored markers represent the density of two strains (green for GFP labeled strain; red for RFP labeled strains) growing in monoculture in 30+ experimental replicates, black lines represent the best-fit model. **(A)** Strain DH5! labeled with GFP, strain TG1 labeled with RFP. Experiment started by diluting stationary phase bacteria into fresh media, yielding a lag phase culture in which lag phase is longer for the green strain. **(B)** Strain DH5! labeled with GFP, strain TG1 labeled with RFP. Bacteria were pre-grown in fresh media for 4 hours before the experiment and then diluted into fresh media, such that there is no observable lag phase. **(C)** Strain JM109 labeled with GFP and, strain K12 MG1655-Δfnr labeled with RFP. Experimental conditions as described in (A).

**Table 1.** Estimated growth parameters. 95% confidence intervals, calculated using bootstrap (1000 samples), are given in parentheses. Min doubling time is the minimal time required to double the population density. Densities are in OD_595_; growth rate in hours^-1^, doubling time and lag duration in hours. See Table S2 for additional parameter estimates.

|  | Experiment A |  | Experiment B |  | Experiment C |  |
| --- | --- | --- | --- | --- | --- | --- |
| Strain | GFP | RFP | GFP | RFP | GFP | RFP |
| Parameter |  |  |  |  |  |  |
| Initial density ($N_0$) | 0.125 | 0.124 | 0.286 | 0.23 | 0.188 | 0.204 |
| Max density ($K$) | 0.528 (0.525, 0.532) | 0.650 (0.643, 0.658) | 0.619 (0.612, 0.625) | 0.628 (0.624, 0.632) | 0.633 (0.627, 0.638) | 0.741 (0.735, 0.746) |
| Max specific growth rate | 0.268 (0.262, 0.275) | 0.376 (0.371, 0.382) | 0.256 (0.251, 0.261) | 0.369 (0.355, 0.384) | 0.228 (0.226, 0.231) | 0.42 (0.391, 0.426) |
| Min doubling time | 2.695 (2.636, 2.77) | 1.844 (1.809, 1.88) | 4.372 (4.269, 4.481) | 2.451 (2.397, 2.506) | 3.117 (3.087, 3.147) | 2.075 (2.035, 2.124) |
| Lag duration | 3.93 (3.82, 4.028) | 1.578 (1.513, 1.64) | 0.004 (0.002, 0.013) | 0.014 (0.002, 0.029) | 0.711 (0.684, 0.749) | 0.039 (0.033, 0.081) |

## b. Estimate competition coefficients

### Competition model

To model growth in a mixed culture, we assume that interactions between the strains are solely due to resource competition. Therefore, all interactions are described by the deceleration of the growth rate of each strain in response to growth of the other strain. We derived a new two-strain Lotka-Volterra competition model^13^ based on resource consumption (see Supporting Text 2):

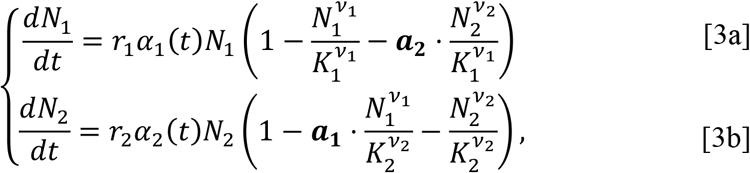

where *N_i_* is the density of strain *i* = 1,2, *r*_*i*_, *K*_*i*_, *ν*_*i*_, *α*_*i*_, *q*_0,*i*_, and *m*_*i*_ are the values of the corresponding parameters for strain *i* (obtained from fitting the growth model (eq. 2) to monoculture growth curve data), and *a*_*i*_ are competition coefficients, the ratios between inter- and intra-strain competitive effects. Note that each strain can have a different limiting resource and resource efficiency, based on the maximum densities *K*_*i*_ and competition coefficients *a*_*i*_ determined for each strain.

### Model fitting

The competition model (eq. 3) was fitted to growth curve data from the mixed culture, in which the combined OD of both strains was recorded over time (i.e. the total density, not the frequency or density of individual strains). The fitting provides estimates for the competition coefficients *a*_*i*_ and was performed by minimizing the squared differences between *N*_1_ + *N*_2_ (the sum of the integrals of the system in eq. 3) and the total OD from the mixed culture (Figure 3A-C). Note that the total density of the mixed culture is usually ignored, despite being easy to produce, and part of the strength of our approach stems from using these data.

## c. Prediction and validation of relative growth

### Model prediction

With estimates of all the competition model parameters, we solved the competition model (eq. 3) using numerical integration, thus providing a prediction for the cell densities *N*_1_(*t*) and *N*_2_(*t*) of the two strains growing in a mixed culture. From these predicted densities, the frequencies of each strain over time were estimated:

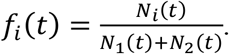

### Experimental validation

The green and red markers and error bars in Figure 3D-F show the results of the competition experiment, that is, the frequency of each strain growing together in a mixed culture. These experimental results are compared to our model predictions in green and red dashed lines and to the exponential model predictions in black dashed lines (see Figure 1 and Introduction for details on the exponential model). Our model performs well and clearly improves upon the exponential model for predicting competition dynamics in a mixed culture: the colored dashed lines match the data much better than the black dashed lines.

**Figure 3.**
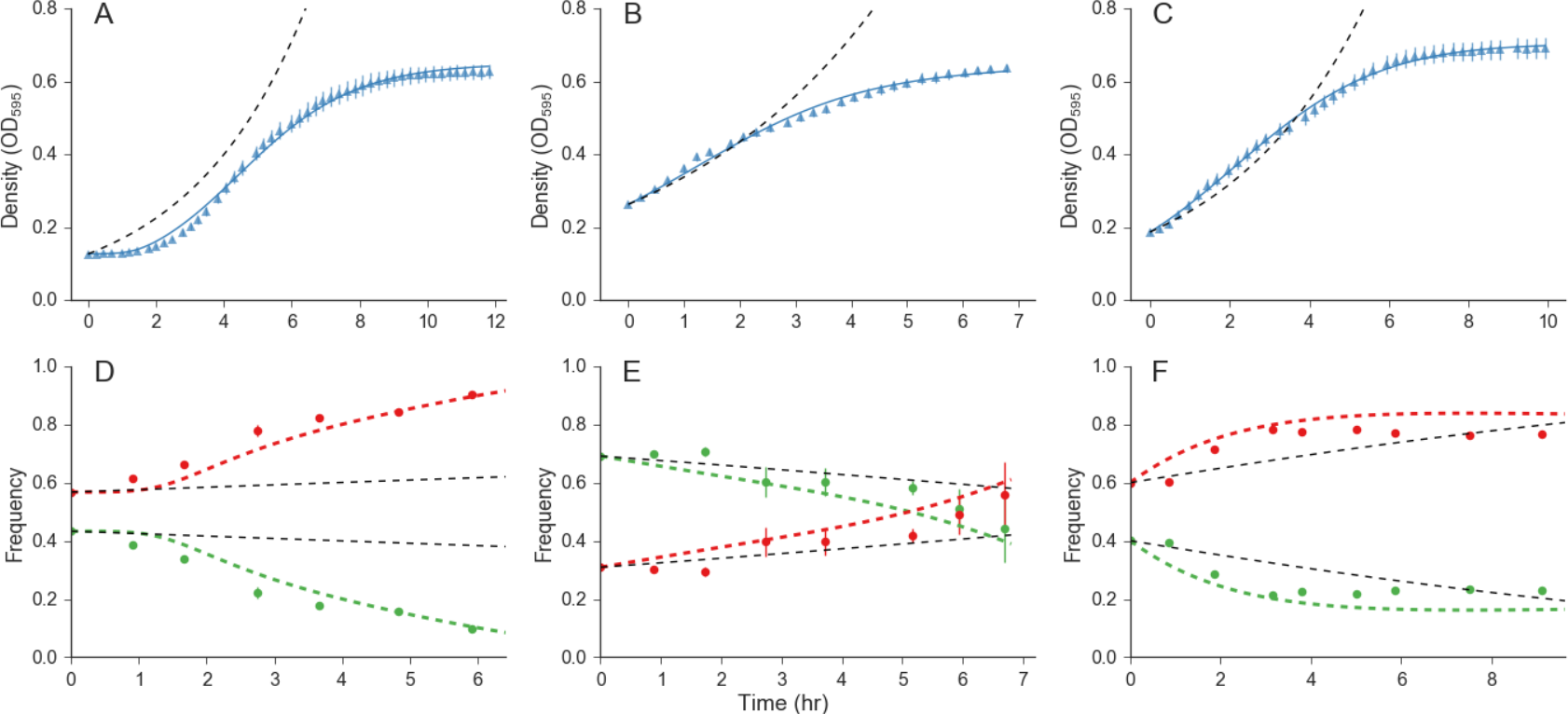
Predicting growth in a mixed culture. Growth of two *E. coli* strains competing for resources in a mixed culture. **(A-C)** Blue triangles show the average overall density in a mixed culture; error bars show standard deviation from 30 replicates (extremely small in B); solid blue lines show the best-fit competition model (eq. 3); dashed black lines show the exponential model prediction (see Figure 1). **(D-F)** Comparison of experimental data (circles) and model prediction (dashed lines; see Figure S3 for confidence intervals) of strain relative frequencies in a mixed culture. Green and red dashed lines show our model predictions; dashed black lines show exponential model predictions. Error bars show standard deviation (hardly seen in D and F). Root mean squared error of model fit (solid blue line to blue triangles A-C) are 0.011 for A, 0.01 for B, and 0.008 for C. Estimated competition coefficients are a_1_=10, a_2_=0.77 for D; a_1_=3.7, a_2_=1.9 for E; a_1_=0.31, a_2_=0.56 for F. Inferred time-averaged selection coefficients are 0. A, 376 for D, 0.182 for E, and 0.124 for F.

## Discussion

We developed a new computational approach to predict relative growth in a mixed culture from growth curves of mono- and mixed cultures, without measuring frequencies of single isolates within the mixed culture. We tested and validated this new approach, which performed far better than the approach commonly used in the literature.

Our approach only assumes that the assayed strains grow in accordance with the growth and competition models: namely, that growth depends on resource availability. Therefore, this approach can be applied to data from a variety of organisms, experiments, and conditions. Growth curve experiments, in which only optical density is measured, require much less effort and resources than pairwise competition experiments, in which the cell frequency or count of each strain must be determined^2,3,8,14^. Current approaches to estimating fitness from growth curves mostly use the growth rate or the maximum population density as a proxy for fitness. However, the growth rate and other proxies for fitness based on a single growth parameter cannot capture the full scope of effects that contribute to differences in overall fitness^15^. In contrast, our new approach integrates several growth phases, allowing a more accurate estimation of relative growth and fitness from growth curve data, and providing information on the specific growth traits that contribute to differences in fitness.

We will release *Curveball*, an open-source software package which implements our new approach (http://curveball.yoavram.com). This software is written in Python^16^ and includes a user interface that does not require prior knowledge in programming. It is free and open, such that additional data formats, growth and competition models, and other analyses can be added by the community to extend its utility.

## Conclusions

We developed and tested a new approach to analyze growth curve data, and applied it to predict growth of individual strains within a mixed culture. This approach can improve fitness estimation from growth curve data, has a clear biological interpretation, and can be used to predict and interpret growth in a mixed culture and results of competition experiments.

## Materials and Methods

### Strains and plasmids

*Escherichia coli* strains used were DH5α (Berman lab, Tel-Aviv University), TG1 (Ron lab, Tel-Aviv University), JM109 (Nir lab, Tel-Aviv University), and K12 MG1655-Δfnr (Ron lab, Tel-Aviv University). Plasmids contain a GFP or RFP gene and genes conferring resistance to kanamycin (Kan^R^) and chloramphenicol (Cap^R^) (Milo lab, Weizmann Institute of Science^17^).

### Media

All experiments were performed in LB media (5 g/L Bacto yeast extract (BD, 212750), 10 g/L Bacto Tryptone (BD, 211705), 10 g/L NaCl (Bio-Lab, 190305), DDW 1 L) with 30 µg/mL kanamycin (Caisson Labs, K003) and 34 µg/mL chloramphenicol (Duchefa Biochemie, C0113). Green or red fluorescence of each strain was confirmed by fluorescence microscopy (Nikon Eclipe Ti, Figure S1).

### Growth and competition experiments

All experiments were performed at 30°C. Strains were inoculated into 3 ml LB+Cap+Kan and grown overnight with shaking. Saturated overnight cultures were diluted into fresh media so that the initial OD was detectable above the OD of media alone (1:1-1:20 dilution rate). In experiments that avoided a lag phase, cultures were pre-grown until the exponential growth phase was reached as determined by OD measurements (4-6 h). Cells were then inoculated into 100 µL LB+Cap+Kan in a 96-well flat-bottom microplate (Costar):

- 32 wells contained a monoculture of the GFP-labeled strain
- 30 wells contained a monoculture of the RFP-labeled strain
- 32 wells containing a mixed culture of both GFP- and RFP-labeled strains
- 2 wells contained only growth medium

The cultures were grown in an automatic microplate reader (Tecan infinite F200 Pro), shaking at 886.9 RPM, until they reached stationary phase. OD_595_ readings were taken every 15 minutes with continuous shaking between readings.

Samples were collected from the incubated microplate at the beginning of the experiment and once an hour for 6-8 hours: 1-10 µL were removed from 4 wells (different wells for each sample), and diluted into cold PBS buffer (DPBS with calcium and magnesium; Biological Industries, 02-020-1). These samples were analyzed with a fluorescent cell sorter (Miltenyi Biotec MACSQuant VYB). GFP was detected using a 488nm/520(50)nm FITC laser. RFP was detected with a 561nm/615(20)nm dsRed laser. Samples were diluted further to eliminate "double" event (events detected as both "green" and "red" due to high cell density) and noise in the cell sorter^2^.

### Data analysis

Fluorescent cell sorter output data was analyzed using R^18^ with the *flowPeaks* package that implements an unsupervised flow cytometry clustering algorithm^19^. Growth curve data were analyzed using *Curveball*, a new open-source software written in Python^16^ that implements the approach presented in this manuscript. The software includes both a programmatic interface (API) and a command line interface (CLI), and therefore does not require programming skills. The source code makes use of several Python packages: NumPy^20^, SciPy^21^, Matplotlib^22^, Pandas^23^, Seaborn^24^, LMFIT^25^, Scikit-learn^26^, and SymPy^27^.

### Fitting growth models

To fit models to data we used the least-squares non-linear curve fitting procedure^21,25^. We then calculate the Bayesian Information Criteria (BIC) of several nested models, defined by fixing some of the parameters (see Supporting Text 1, Figure S2, and Table S1). BIC is given by:

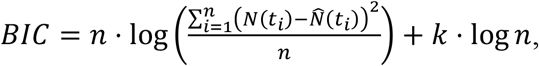

where *k* is the number of model parameters, *n* is the number of data points, *t*_1_ are the time points, *N*(*t*_1_) is the optical density at time point *t*_1_, and 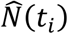 is the expected density at time point *t*_*i*_ according to the model. We selected the model with the lowest BIC^28,29^. Other metrics for model selection can be used with *Curveball*, but BIC was chosen for its simplicity and flexibility.

### Fitting exponential models

The following represents a common approach for estimating growth rates from growth curve data, and was used as a benchmark for our new approach (see Figure 1 and black dashed lines in Figure 3). A polynomial is fitted to the mean of the growth curve data *N(t)*. The time of maximum growth rate *t_max_* is found by differentiating the fitted polynomial and finding the maximum of the derivative. Values *a* and *b* are found such that *f(t)=b+at* describes a tangent line at the point of maximum growth (*t_max_, N(t_max_)*). The intercept *b* and the slope *a* are interpreted as the initial density *N_0_=e^b^* and the growth rate *r=a* in an exponential growth model *N(t)=N_0_e^rt^* (*N_0_* is usually disregarded).

### Selection coefficients estimation

Selection coefficients were estimated from pairwise competition results using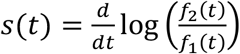 where *f*_1_(*t*) and *f*_2_(*t*) are the predicted frequencies of the strains and *t* is time^1^. The resulting *s_t_* values were then averaged across time. Note that these estimates can depend on the experimental conditions, such as duration, media, temperature, and strain composition.

### Data availability

Data deposited on *figshare* (doi:10.6084/m9.figshare.3485984).

### Code availability

Source code will be available upon publication at https://github.com/yoavram/curveball; an installation guide, tutorial, and documentation will be available upon publication at http://curveball.yoavram.com.

### Figure reproduction

Data was analyzed and figures were produced using a Jupyter Notebook^30^ that will be available as a supporting file and at https://github.com/yoavram/curveball_ms.

## Acknowledgments

We thank Y. Pilpel, D. Hizi, I. Françoise, I. Frumkin, O. Dahan, A. Yona, T. Pupko, A. Eldar, I. Ben-Zion, E. Even-Tov, H. Acar, J. Friedman, J. Masel, M.W. Feldman, and E. Rosenberg, for helpful discussions and comments, and L. Zelcbuch, N. Wertheimer, A. Rosenberg, A. Zisman, F. Yang, E. Shtifman Segal, I. Melamed-Havin, and R. Yaari for sharing materials and experimental advice. This research has been supported in part by the Israel Science Foundation 1568/13 (LH) and 340/13 (JB), the Minerva Center for Lab Evolution (LH), Manna Center Program for Food Safety & Security, the Israeli Ministry of Science & Technology, and Stanford Center for Computational, Evolutionary and Human Genomics (YR), TAU Global Research and Training Fellowship in Medical and Life Science and the Naomi Foundation (MB), and the European Research Council (FP7/2007-2013)/ERC grant 340087 (JB).

## Author contributions

All authors designed the experiments, analyzed data, discussed the results and edited the manuscript. Y.R. and L.H. developed the model and wrote the manuscript. U.O. advised on statistical analysis. Y.R. wrote the source code. Y.R., E.D.G. and M.B. performed the experiments. M.B. performed fluorescent microscopy. J.B. advised and gave support to all experiments. L.H. supervised all the work.

## Supporting material

### Supporting Text 1: Monoculture model

We derive our growth models from a resource consumption perspective^13,31^. We denote by *R* the density of a limiting resource, and by *N* the density of the cell population, both in total mass per unit of volume.

We assume that the culture is well-mixed and homogeneous and that the resource is depleted by the growing cell population without being replenished. Therefore, the intake of resources occurs when cells meet resource via a mass action law with resource intake rate ℎ. Once inside the cell, resources are converted to cell mass at a conversion rate of ∊. Cell growth is assumed to be proportional to *R* ⋅ *N*, whereas resource intake is proportional to a power of cell density, *R* ⋅ *N*^*ν*^. We denote *Y* ∶= *N*^*ν*^.

We can describe this process with differential equations for *R* and *N*:

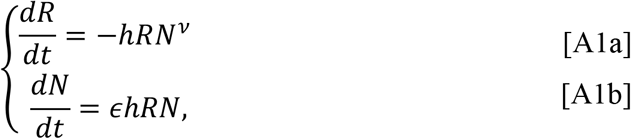

These equations can be converted to equations in *R* and *Y*:

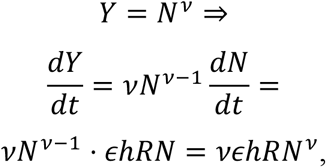

which yields

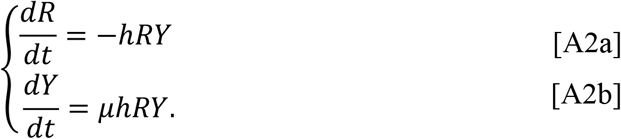

with *μ* = *∊ν*.

To solve this system, we use a conservation law approach by setting *M* = *μR* + *Y*^32^. We find that *M* is constant

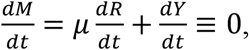

and we can substitute *μR* = *M* − *Y* in eq. A2b to get

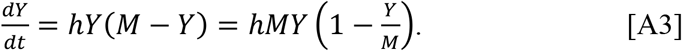

Substituting again 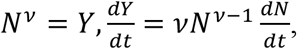 and defining 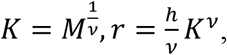 we get

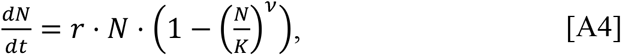

which is the Richards differential equation^33^, with the maximum population density *K* and the specific growth rate in low density *r*. To the best of our knowledge this the first derivation of the Richards differential equation from a resource consumption perspective.

We solve eq. A4 via eq. A3, which is a logistic equation and therefore has a known solution. Setting the initial cell density *N* 0 = *N*_0_ we have

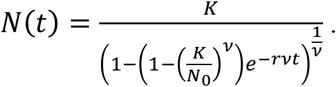

Eq. A4 is an autonomous differential equation (*dN*/*dt* doesn’t explicitly depend on *t*). To include a lag phase, Baranyi and Roberts^11^ suggested to add an adjustment function *α*(*t*), which makes the equation non-autonomous (explicitly dependent on *t*):

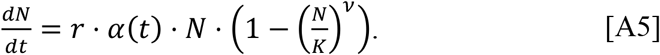

Baranyi and Roberts suggested a Michaelis-Menten type of function^34^

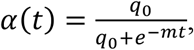

which has two parameters: *q_0_* is the initial physiological state of the population, and *m* is rate at which the physiological state adjusts to growth conditions. Integrating *α*(*t*) gives

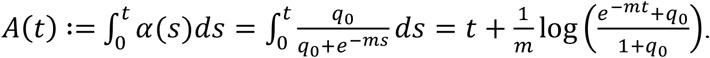

Therefore, integrating eq. A5 produces eq. 2.

The term 1 − (*N*/*K*)^*ν*^ is used to describe the deceleration in the growth of the population as it approaches the maximum density *K*. When *ν* = 1, the deceleration is the same as in the standard logistic model 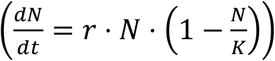 and the density at the time of the maximum population growth 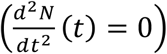 is half the maximum density, 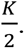 When *ν* > 1 or 1 > *ν*, the deceleration is slower or faster, respectively, and the density at the time of the maximum growth rate is *K*/(1 + *ν*)^1/*ν*^ (Richards 1959, substituting W = N, A = K, m = ν + 1, k = r ⋅ ν).

We use six forms of the Baranyi-Roberts model (Figure S2, Table S1). The full model is described by eq. 2 and has six parameters. A five-parameter form of the model assumes *ν* = 1, as in the standard logistic model, but still incorporates the adjustment function *α*(*t*) and therefore includes a lag phase. Another five-parameter form has both rate parameters set to the same value (*m* = *r*), which was suggested to make the fitting procedure more stable^34,35^. A four-parameter form has both of the previous constraints, setting *m* = *r* and *ν* = 1^34^. Another four-parameter form of the model has no lag phase, with 1/*m* = 0 ⇒ *α*(*t*) ≡1, which yields the Richards model^33^, also called the θ-logistic model^36^, or the generalized logistic model. This form of the model is useful in cases where there is no observed lag phase: either because the population adjusts very rapidly or because it is already adjusted prior to the growth experiment, possibly by pre-growing it in fresh media before the beginning of the experiment. The last form is the standard logistic model, in which *ν* = 1 and 1 *m* = 0.

### Supporting Text 2: Mixed culture model

We consider the case in which two species or strains grow in the same culture, competing for a single limiting resource, similarly to eq. A1:

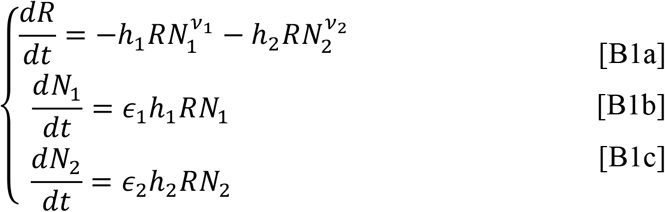

We define 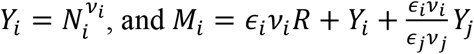 (where *j* is 1 when *i* is 2 and vice versa) to find that 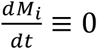 and *M*_*i*_ is constant. We then substitute 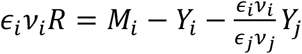 into the differential equations for 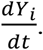 Denoting 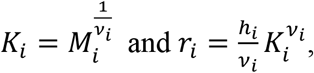 we get

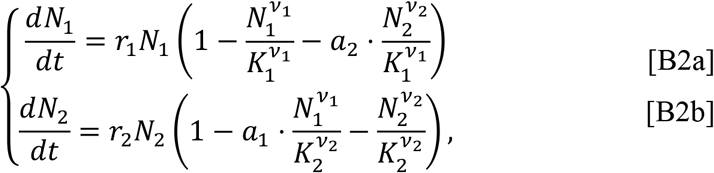

where 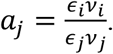

We get a similar result if each strain is limited by a different resource that both strains consume:

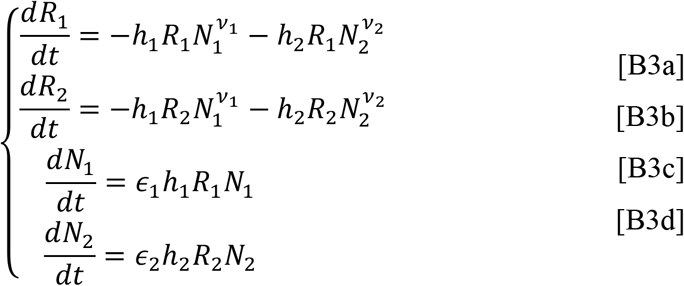

Here, we notice first that 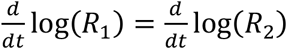 and therefore 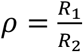 is a constant. We then substitute *R*_1_ = *R, R*_2_ = *ρR* in eqs. B3 and continue as above. This changes the definition of 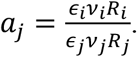

If the intake rates depend only on the resource then

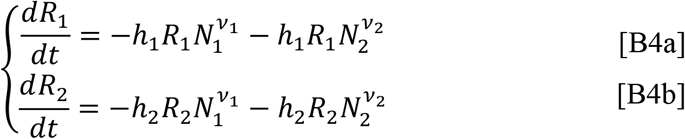

Then we define 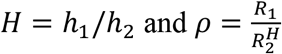 and again continue as above.

### Supporting figures

**Figure S1.**
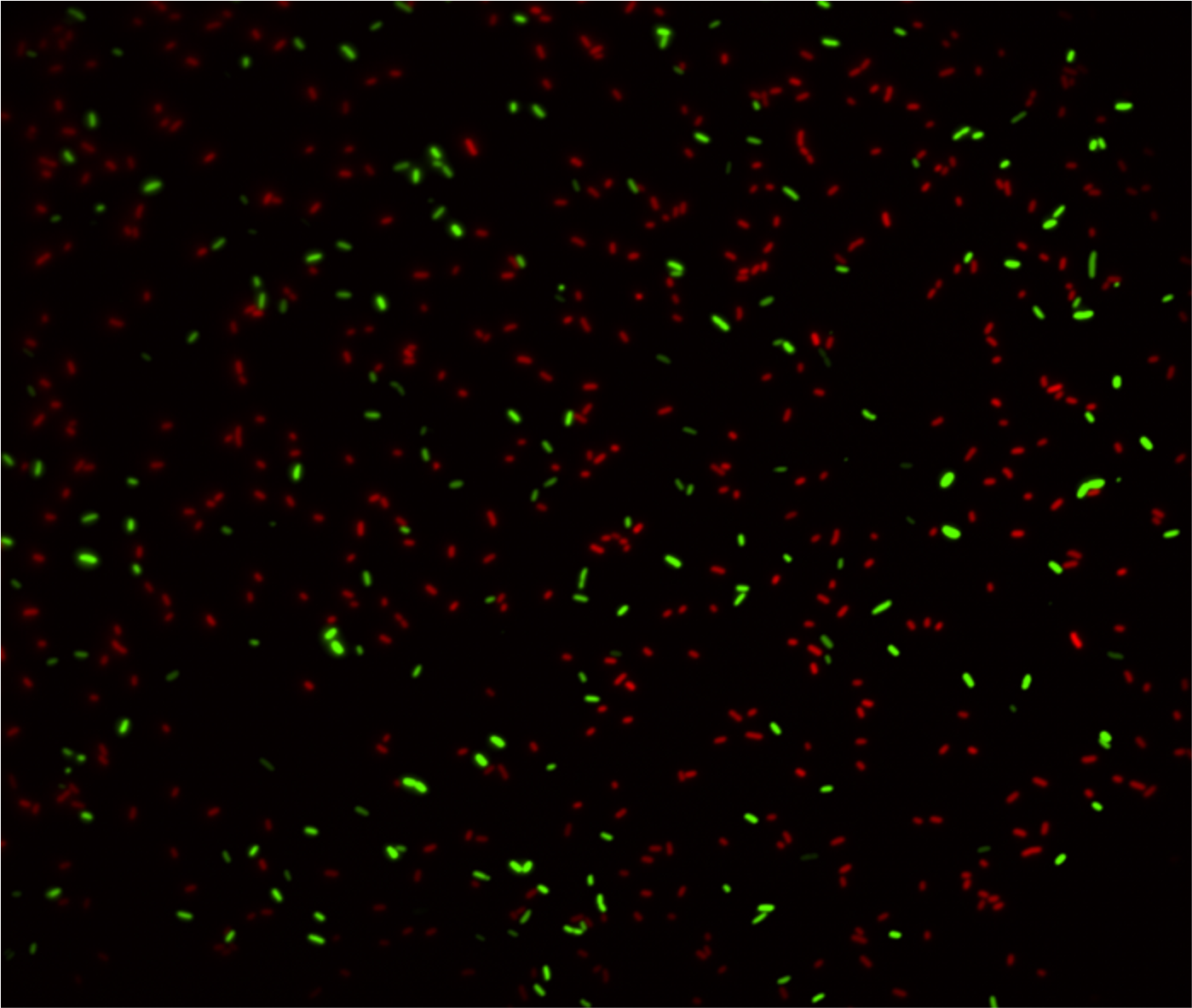
Fluorescence microscopy of *E. coli* strains carrying GFP or RFP. Image of a mixture of DH5α-GFP and TG1-RFP cells.

**Figure S2.**
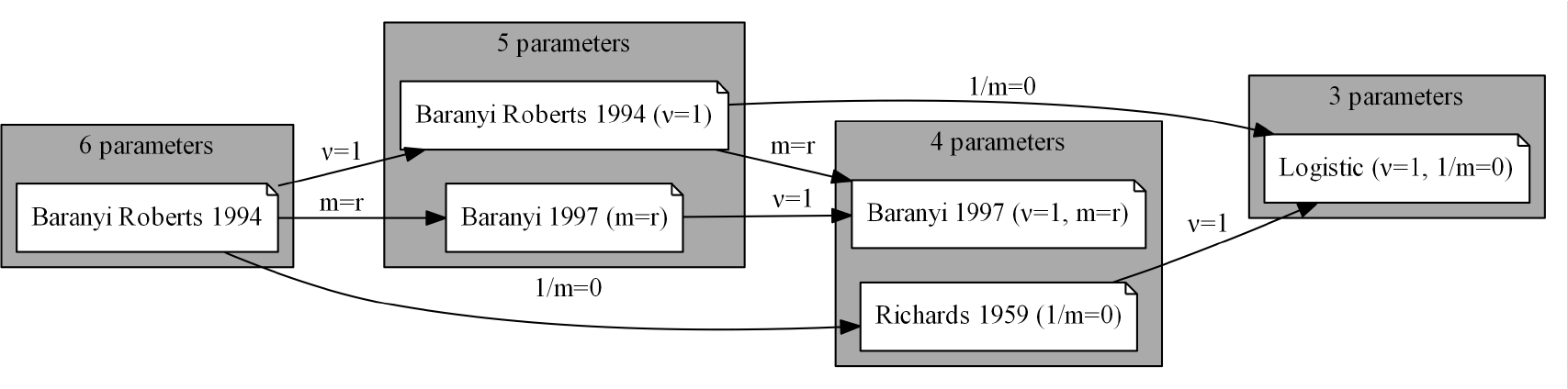
Growth models hierarchy. The Baranyi-Roberts model and five nested models defined by fixing one or two parameters. See Supporting Text 1 and Table S1 for more details.

**Figure S3.**
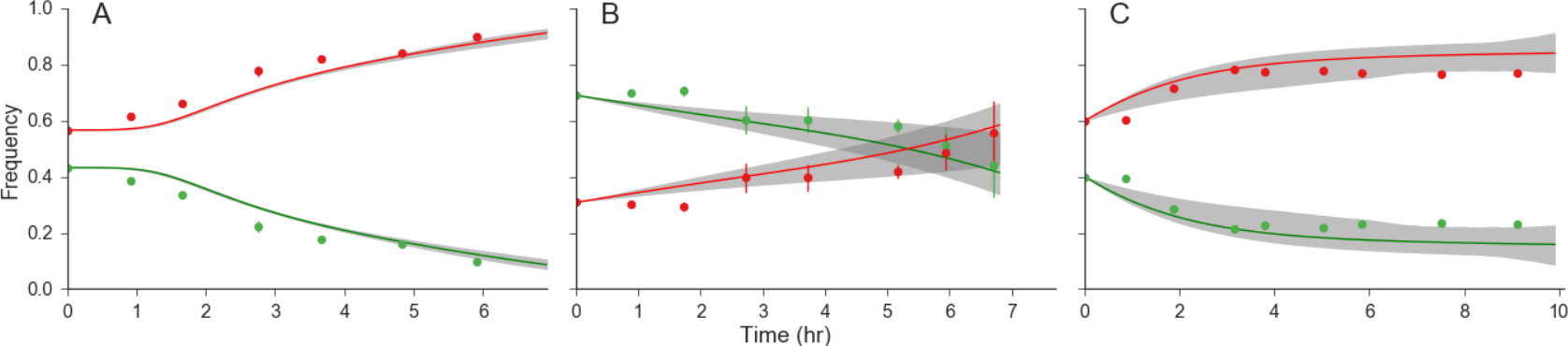
Mixed culture growth predictions with confidence intervals. The green and red lines and markers correspond to the dashed green and red lines and the markers in Figure 3D-F, respectively. The gray area shows the 95% confidence interval, calculated using bootstrap (1000 samples).

### Supporting tables

**Table S1.** Growth models. The table lists the growth models used for fitting growth curve data. All models are defined by eqs. 1 and 2, by fixing specific parameters. ***N*_0_** is the initial population density; ***K*** is the maximum population density; ***r*** is the specific growth rate in low density; ***v*** is the surface to mass ratio; ***q*_0_** is the initial physiological state; ***m*** is the physiological adjustment rate. Note that when **1/*m* = 0**, the value of ***q*_0_** is irrelevant. See also the hierarchy diagram in Figure S2 and a detailed discussion in Supporting text 1.

| Model name | # Parameters | Free Parameters | Fixed Parameters | References |
| --- | --- | --- | --- | --- |
| **Baranyi Roberts 1994** | 6 | $N_0, K, r, v, q_0, m$ | - | 11 |
| **Baranyi 1997** | 5 | $N_0, K, r, v, q_0$ | $m = r$ | - |
| **Baranyi Roberts 1994** | 5 | $N_0, K, r, q_0, m$ | $v = 1$ | - |
| **Richards 1959** | 4 | $N_0, K, r, v$ | $\frac{1}{q_0} = \frac{1}{m} = 0$ | 33 |
| **Baranyi 1997** | 4 | $N_0, K, r, q_0$ | $v = 1$<br>$m = r$ | 34 |
| **Logistic** | 3 | $N_0, K, r$ | $v = 1$<br>$\frac{1}{q_0} = \frac{1}{m} = 0$ | 37 |

**Table S2.** Estimated parameters from growth model fitting. * denotes fixed parameters; - denotes invalid parameter values.

|  | Experiment A |  | Experiment B |  | Experiment C |  |
| --- | --- | --- | --- | --- | --- | --- |
| *Strain* | GFP | RFP | GFP | RFP | GFP | RFP |
| *Parameter* |  |  |  |  |  |  |
| $N_0$ | 0.125 | 0.124 | 0.286 | 0.23 | 0.188 | 0.204 |
| $K$ | 0.528 | 0.65 | 0.619 | 0.628 | 0.633 | 0.741 |
| $r$ | 0.376 | 0.587 | 0.304 | 0.484 | 8 | 8 |
| $v$ | 2.636 | 1* | 2.484 | 1.491 | 1* | 0.164 |
| $q_0$ | 0.032 | 0.008 | -* | -* | 0.039 | 0.393 |
| $v$ | 0.937 | 3.735 | -* | -* | 0.188 | 0.104 |

